# The First Chromosome-Scale Genome Assembly for *Dermacentor reticulatus*: A Key Vector of Tick-Borne Pathogens of Public and Veterinary Health Importance in Europe

**DOI:** 10.1101/2025.10.09.681344

**Authors:** Nina Billows, Mia L. White, Joseph Thorpe, Matthew Higgins, Emma Collins, Myrela Santos de Jesus, Mojca Kristan, Steven Pullan, Sarah M. Biddlecombe, Faye V. Brown, Taane G. Clark, Kayleigh M. Hansford, Jolyon M. Medlock, Susana Campino

## Abstract

**Background:** *Dermacentor reticulatus* is a key tick species across Europe and an established vector of multiple pathogens that affect both human and animal health. Despite its significant role in disease transmission, genomic data for this species remain limited. Here, we present the first chromosome-scale genome assembly of *D. reticulatus*, constructed using Oxford Nanopore long-read sequencing data.

**Methods:** High molecular weight DNA was extracted from a female *Dermacentor reticulatus* collected in Devon, UK, and sequenced using Oxford Nanopore long-read technology. Genome assemblies were generated using both guided and unguided approaches, followed by repeat masking, gene prediction, and functional annotation. Genome completeness was assessed using BUSCO, and comparative, phylogenetic, and functional analyses were performed against other *Dermacentor* and tick species.

**Results:** This chromosome-scale genome assembly revealed a repeat-rich genome, with approximately 61.9% of the total sequence consisting of repetitive elements. Benchmarking universal single-copy ortholog (BUSCO) analysis demonstrated strong genome completeness, with guided assembly (chromosome) achieving a score of 97.1%, closely matching those of related *Dermacentor* species. For comparison, unguided assembly (scaffold) yielded a BUSCO score of 96.7%. Gene annotation following repeat masking resulted in BUSCO completeness scores of 95.1% (guided) and 95.0% (unguided). Functional characterisation included Pfam domain assignment and Gene Ontology analysis. Additionally, we assembled the mitochondrial genome (15,103 bp), comprising 38 genes, providing further insight into *D. reticulatus* phylogenetic placement.

**Conclusions:** This genomic resource establishes a foundation for functional genomics, evolutionary studies and supports future research in vector biology and the control of tick-borne diseases.

## Background

*Dermacentor reticulatus*, also known as the ornate cow tick or European meadow tick, is a vector of medical and veterinary importance, capable of transmitting a broad range of pathogens. Predominantly found across Europe and parts of Western Asia, *D. reticulatus* thrives in a variety of habitats with preference for grasslands, pastures, meadows, sand dunes, marshy areas and grassy paths in woodlands. This three-host species commonly feeds on small, medium and -large mammals such as small mammals, rabbits, deer, horses, and livestock, while also opportunistically biting humans and companion animals, especially dogs [1-4]. In the UK, *D. reticulatus* has mostly been observed across coastal regions of Southwest England and West Wales associated with sand dune rabbit burrow systems and coastal maritime grassland, as well as urban green spaces in Essex [5]. Historically, the distribution of *D. reticulatus* was confined to regions with milder climates [6]. However, this species has expanded into cooler and more northerly regions, including Belgium, the Netherlands, Germany, Poland, Hungary, Slovakia and new foci in south-east England [3, 5, 6] and is essentially known as a winter tick.

This trend is mostly associated with movement of infested livestock but could be driven by other anthropogenic factors [5, 7, 8]. The wide geographic distribution of *D. reticulatus* raises public and animal health concerns, as this tick is a known vector for some pathogens (e.g., *Babesia, Anaplasma, Rickettsia, Theileria*) and has been found to carry other agents, including *Bartonella, Francisella, Coxiella*, Omsk haemorrhagic fever virus and tick-borne encephalitis virus, though vector competence for these remains unconfirmed [9, 10]. Surveillance of *D. reticulatus* is therefore essential for early detection of range shifts and associated disease threats. While the morphology and some elements of the ecology of *D. reticulatus* are well characterised, including its high reproductive rate, flexible habitat preferences, and ability to transmit diverse pathogens, there remains a significant genomic knowledge gap [10]. This gap limits molecular-level understanding of its biology and vector capacity.

Currently, no chromosome-scale reference genome exists for *D. reticulatus*. Genomic resources are critical for accurate species identification, detection of cryptic population structure and monitoring of pathogen carriage. Moreover, they enable population genomics and comparative analyses to uncover adaptive traits linked to vector competence, climate tolerance or ecological plasticity. The use of microsatellite loci and mitochondrial sequence data have provided some insight into *D. reticulatus* ancestral admixture [11]. However, further whole genome sequencing studies are invaluable for enhancing tick-borne disease surveillance and informing the development of novel vector control strategies, with the potential for reducing the health risks posed by this expanding vector.

To date, reference genomes with varying quality have been generated for related *Dermacentor* species largely distributed across Asia and North America [11,12]. For instance, *D. silvarum* (GCA_013339745.2), a dominant tick species in Northeast China, was sequenced using an integrated approach combining Illumina, PacBio, and Hi-C technologies, enabling chromosome-level resolution [12]. Three additional ticks belonging to the *Dermacentor* genus have chromosome-scale genome assemblies available, including *D. albipictus* (GCA_038994185.2), *D. andersoni* (GCA_023375885.3) and *D. variabilis*, all found within North America and Canada, whilst a scaffold-level is available for *D. reticulatus* (GCA_051549955.1) [13, 14]. These genomic data provide a robust framework for comparative studies, but do not adequately represent European tick populations.

In response to the expanding distribution of *D. reticulatus* and the rising incidence of tick-borne diseases in Europe, we identified an urgent need for a reference genome for this species [10]. In this study, we present the first chromosome-scale genome assembly for *D. reticulatus*, generated from a female tick collected in Devon, England (March 2024), using Oxford Nanopore Technology (ONT) long-read sequencing. We evaluate assembly and annotation quality, conduct comparative analyses with other *Dermacentor* genomes, and highlight the genomic architecture and functional gene content associated with its hematophagous lifestyle. This assembly provides a foundational genomic resource to support future surveillance, population genomics, and functional studies aimed at better understanding the biology, adaptation, and control of this medically and veterinary important tick species.

## Methods

### Sample Collection and Sequencing

A single *D. reticulatus* female sample was collected in a known established area in Devon, England using flagging. The sample as homogenised using a pestle and DNA was extracted using Dynabeads™ SILANE Genomic DNA Kit using the manufacturer’s instructions, to produce high genomic weight DNA. The concentration of the DNA was assessed using the Qubit Fluorometer with the dsDNA high sensitivity kit (Invitrogen). The 4200 TapeStation system, with Genomic DNA ScreenTape assay (Agilent), was used to assess the quality and quantity of DNA, and the sample with the highest molecular weight DNA was selected (80% DNA with peak at 20,000bp).

A DNA library of the *D. reticulatus* was prepared using the Ligation Sequencing Kit V14 (SQK-LSK114; Oxford Nanopore Technologies (ONT), UK), following the manufacturer’s protocol. The DNA library was sequenced on an R10.4.1 PromethION flow cell (FLO-PRO114M; ONT, UK). Raw reads were basecalled using Dorado v0.9.1 (super high accuracy model) to convert the sequence signals (“squiggle”) with –trim to remove adaptor sequences.

### Genome Assembly

The *D. reticulatus* genome was assembled using an established pipeline (Available at: https://github.com/MatthewHiggins2017/DenovoAssemblyTools/). Kraken2 v2.1.3 was used to assign taxonomic labels to raw ONT reads and generate a contamination report [15]. No eukaryotic contaminants were reported and raw reads were subsequently filtered to remove sequences matching ‘Archaea’ in the Kraken2 report using KrakenTools (extract_kraken_reads.py) [16]. The remaining reads were scrubbed using yacrd v1.0.0 to exclude any poor quality or chimeric sequences [17]. Reads were filtered further using chopper v0.6.0 to extract reads with a minimum length of 1000 and minimum quality of 12 [18]. The high-quality reads underwent unguided reference assembly using Flye v2.9.2 [19]. Several read error scores were trialled (0.04, 0.06 and 0.08). A read error of 0.06 generated the highest BUSCO score so was used for downstream analysis. The unguided scaffolds underwent additional guided scaffolding using the available *D. silvarum* genome (GCA_013339745.2) [20]. This genome was chosen as the reference guide due to its high-quality, close phylogenetic relationship with *D. reticulatus* as determined by mtDNA analysis (compared with other *Dermacentor* species), and expected chromosome number (previously determined by karyotyping: 10 Autosomal Chromosomes, 1 Sex Chromosome) [21]. Guided scaffolding was carried out using NtLink v1.3.9 with a maximum gap parameter of 500 and gap-fill option [22]. The guided genome assembly was improved using RagTag v2.1.0 and remaining gaps were filled using TGS-GapCloser v1.2.1 [23, 24]. Finally, the unguided and guided genome assemblies were polished using racon v1.5.0, before screening for contaminants and adaptors using the NCBI Foreign Contamination Screen (FCS) suite [25, 26].

### Genome Assembly Quality Assessment

The quality of genome assemblies were assessed using BUSCO v5.8.2 (-m genome -l arachnida_odb10) after each stage of the pipeline to assess the impact of each process on quality [27]. Final quality assessments were conducted for the unguided and guided *D. reticulatus* genome assemblies and benchmarked against assembly metrics from other available *Dermacentor* genomes in the GenBank database, including *D. silvarum* (GCA_013339745.2, 11 chromosomes), *D. albipictus* (GCA_038994185.2, 10 chromosomes) and *D. andersoni* (GCA_023375885.3, 11 chromosomes) [20]. Approximate genome size was estimate using Jellyfish v2.3.1 (21 k-mers) [28].

### Repeat Masking and Genome Annotation

Before undergoing annotation (and guided scaffolding), it is important to mask repeats to ensure they do not affect downstream annotation. RepeatMasker v4.1.7 and the Repbase database (Eukaryota and Arthropoda, 2018-10-26) were first used to identify and classify repeats in the *D. reticulatus* genome [29, 30]. This yielded a small percentage of repeats, prompting further *de novo* modelling of repeats using RepeatModeler v 2.0.1 [31]. The results from both outputs were combined to hard and soft-mask sequences using RepeatMasker. The soft masked sequences were used in downstream annotation. Several approaches were trialled and compared to identify the best tool for genome annotation. In the absence of RNA-seq data, tools that annotate the genome assemblies using homology and *ab initio* approaches were run. This included lift-off v1.6.3, BRAKER v3.0.8 and GALBA v1.0.11 [32-34]. For the *ab initio* approaches GALBA yielded the best results and was prioritised over BRAKER for forming the final annotation. GALBA utilised protein sequences of closely related *Dermacentor sp* to generate a training gene set for AUGUSTUS software during the annotation process. Additionally, tRNA sequences were also annotated using tRNAscan-SE. The quality of genome annotation was assessed using BUSCO v5.8.2 (-m protein -l arachnida_odb10) and was compared between the unguided and guided *D. reticulatus* assemblies and aforementioned *Dermacentor sp*. assemblies [27].

### Nucleotide and Gene Synteny

To further investigate potential misassemblies in the *D. reticulatus* assemblies, nucleotide-level synteny analysis was performed against the *D. silvarum* genome. Whole genome alignment was performed using minimap2 v2.28 (-ax asm5) between the *D. silvarum* reference and masked *D. reticulatus* assemblies [35]. Mapped regions were filtered to retain high-quality matches (MAPQ>60) to exclude any potential unmasked repeats, and were used to create a ‘links’ file. The links between the *D. reticulatus* assembly and *D. silvarum* were visualised using the PyCircos v3.10.0 package to interpret nucleotide-level synteny (https://github.com/ponnhide/pyCircos). In addition, synteny was also explored at the gene level. Coding sequences for *D. reticulatus* and *Dermacentor sp*. assemblies were extracted, and inputted into jcvi and MCScan (Python version) for pairwise synteny searches and macrosynteny visualisation [36].

### Phylogenetic Orthology Inference

Analysis of orthologues was conducted using the OrthoFinder platform v2.5.5 [37]. Amino acid sequence files (*.faa) for the unguided and guided *D. reticulatus* assemblies were compared to other scaffold- or chromosome-level tick assemblies, including *D. albipictus* (GCF_038994185.2), *D. andersoni* (GCA_023375885.3), *D silvarum* (GCF_013339745.2), *Haemaphysalis longicornis* (GCA_013339765.2), *Ixodes scapularis* (GCF_016920785.2), *Ixodes persulcatus* (GCA_013358835.2), *Ornithodoros turicata* (GCF_037126465.1), *Rhipicephalus microplus* (GCF_013339725.1) and *Rhipicephalus sanguineus* (GCF_013339695.2) [20, 38]. A tree based on orthogroups was built using the platform (-M msa [Default = mafft], -T iqtree). The resulting comparative genomics statistics were used for downstream analyses, and the tree was visualised using iTOL (https://itol.embl.de/). [39].

### Functional Annotation of Protein Sequences

Annotated sequences were extracted and InterProScan (5.73-104.0) was used to predict domains, motifs, and functional annotations associated within the *D. reticulatus* genome using inbuilt databases (E-value<1E-30) [40]. To gain a more comprehensive understanding of the domains and their differences between groups, we searched for the abundance of each Pfam ID within a set of pre-defined groups. These included (1) the *Dermacentor* genus, (2) other aforementioned tick species, (3) blood-feeding insects (*Aedes aegypti:* GCF_002204515.2, *Anopheles gambiae:* GCF_943734735.2 and *Aedes albopictus*: GCF_035046485.1) and (4) non-blood-feeding arachnids (*Centruroides sculpturatus*: GCF_000671375.1 and *Parasteatoda tepidariorum*: GCF_043381705.1) [20, 41-44]. We first calculated the number of each Pfam domain ID and calculated the fold change *Dermacentor* and all other groups. If the median fold-change was >2, the domain ID was mapped to corresponding GO terms. The median fold change was subsequently estimated for each GO term and input into REVIGO for enhanced visualisation and interpretation [45]. Domains of interest associated with key functions for tick biology were identified and predicted *D. reticulatus* amino acid sequences were identified (E-value<1E-10) (**Supplementary Table S12**). The *D. reticulatus* amino acid sequences were also blasted against the *Dermanyssus gallinae* transcriptome to predict the presence or absence of proteins involved in the heme biosynthesis pathway [46].

### Mitochondria Assembly and Annotation

The mitochondrial genome was assembled separately from the nuclear genome. Firstly, mitochondrial amino acid sequences were extracted from the *D. silvarum* (GCA_013339745.2) assembly and used to make a DIAMOND v2.1.11 database [47]. The high-quality filtered *D. reticulatus* reads were blasted against this database and filtered to obtain reads that significantly matched the mitochondria reference (E-value <1e^-5^). The remaining reads were sorted and deduplicated using seqtk before undergoing unguided scaffolding using Flye v2.9.2 (--genome-size 15k) [19]. This yielded a single, circular contig which underwent annotation using MitoZ v3.6 [48].

### Ixodida Mitochondria Tree Analysis

All Ixodida mitochondria sequences were downloaded from the NCBI Organelle database (https://www.ncbi.nlm.nih.gov/datasets/organelle/, Accessed: March 2025) and aligned alongside the *D. reticulatus* mitochondria sequence using MAFFT v7.525 [49]. A maximum-likelihood tree was built using IQ-TREE v 2.3.6 with 1,000 bootstraps and automated model finder selection (-B 1000 -m MFP -T AUTO). The mitochondria tree was visualised using iTOL (https://itol.embl.de/) [39, 50].

## Results

### Assessment of Genome Assembly Quality and Statistics

Overall, ONT sequencing yielded good quality unguided and guided *D. reticulatus* assemblies that are comparable to those for other *Dermacentor* species (**Table 1**). The estimated genome size of *D. reticulatus* was 2.3 Gb, consistent with values reported for other species. The initial overall number of scaffolds was significantly greater for *D. reticulatus* (unguided: 33,814 and guided: 21,637), when compared to *D. silvarum* (N=1,665), *D. andersoni* (N=791) and *D. albipictus* (N=568). The use of *D. silvarum* to guide scaffolding reduced the number of scaffolds by 12,177 and facilitated the formation of putative chromosomes. *D. silvarum* was selected because it is the most closely related species to *D. reticulatus*, supported by phylogenomic analysis of mitochondrial sequences and has a publicly available high-quality, chromosome-level genome assembly [51]. A total of 11 putative chromosomes were formed, matching the chromosome numbers reported for *D. silvarum* and *D. andersoni*, and aligning with previous karyotyping studies of *Dermacentor* species [21]. However, only 10 chromosomes are reported for *D. albipictus*, possibly reflecting either a genuine biological difference or methodological variation in assembly.

**Table 1.** Quality Assessment and BUSCO scores (arachnida_odb10) of unguided and guided *Dermacentor reticulatus* assemblies in comparison to *Dermacentor sp*.

|  | *Dermacentor silvarum*<sup>a</sup> | *Dermacentor andersoni*<sup>b</sup> | *Dermacentor albipictus*<sup>c</sup> | *Dermacentor reticulatus* (Unguided) | *Dermacentor reticulatus* (Guided) |
| --- | --- | --- | --- | --- | --- |
| **Assembly Statistics:** |  |  |  |  |  |
| **Number of scaffolds** | 1665 | 791 | 568 | 33814 | 21637 |
| **Number of contigs** | 15182 | 836 | 679 | 34008 | 32969 |
| **Percent gaps** | 0.09% | 0.04% | 0.02% | <0.01% | 0.05% |
| **Scaffold N50** | 189 MB | 205 MB | 181 MB | 463 KB | 185 MB |
| **Contigs N50** | 339 KB | 66 MB | 30 MB | 453 KB | 492KB |
| **Putative number of chromosomes** | 11 | 11 | 10 | - | 11 |
| **Genome size** | 2.5 GB | 2.4 GB | 2.2 GB | 2.3 GB | 2.3 GB |
| **GC content** | 47% | 46.5% | 46.5% | 46.4% | 46.4% |
| **BUSCO Score:** | 96.8% | 97.0% | 97.1% | 96.7% | 97.1% |
| **Complete BUSCOs (C)** | 2841 | 2847 | 2849 | 2837 | 2848 |
| **Complete and single-copy BUSCOs (S)** | 2687 | 2782 | 2742 | 2728 | 2738 |
| **Complete and duplicated BUSCOs (D)** | 154 | 65 | 107 | 108 | 110 |
| **Fragmented BUSCOs (F)** | 42 | 31 | 35 | 45 | 36 |
| **Missing BUSCOs (M)** | 51 | 56 | 50 | 52 | 50 |

<sup>a</sup>*D. silvarum* (GCA\_013339745.2, Jia et al., 2020), <sup>b</sup>*D. andersoni* (GCA\_023375885.3) and <sup>c</sup>*D. albipictus* (GCA\_038994185.2).

The 11 putative chromosomes in the *D. reticulatus* genome ranged in size from 107 to 415 Mb, with chromosome 1 being the largest. Chromosomes were labelled by descending size, following standard convention. The guided scaffolding approach introduced a marginal number of gaps into the assembly in favour of increased contiguity (unguided: <0.01% and guided: 0.05%). GC content was also comparable across genome assemblies for all species (∼46.4%). Combined, this demonstrates the utility of ONT sequence for rapid assembly of tick species with adequate quality, as demonstrated by previous studies [52].

The BUSCO (Benchmarking Universal Single-Copy Orthologs) scores of *D. reticulatus* assemblies (unguided: 96.7% and guided: 97.1%) were comparable to *D. silvarum* (96.8%), *D. andersoni* (97.0%) and *D. albipictus* (97.1%). Guided scaffolding improved the BUSCO score by 0.4%. The number of complete and single copy genes was also high across all *Dermacentor* assemblies (>2800), representing the completeness of each genome and the quality of assembly. The lower number of duplicated, fragmented or missing BUSCOs suggests the genomes are well-assembled and are free from misassemblies, duplications or gaps within these regions. Any potential misassemblies were also identified using nucleotide-level synteny visualisation (**Figure 1**). The repeat-masked unguided and guided assemblies demonstrated strong synteny with the *D. silvarum* genome.

**Figure 1.**
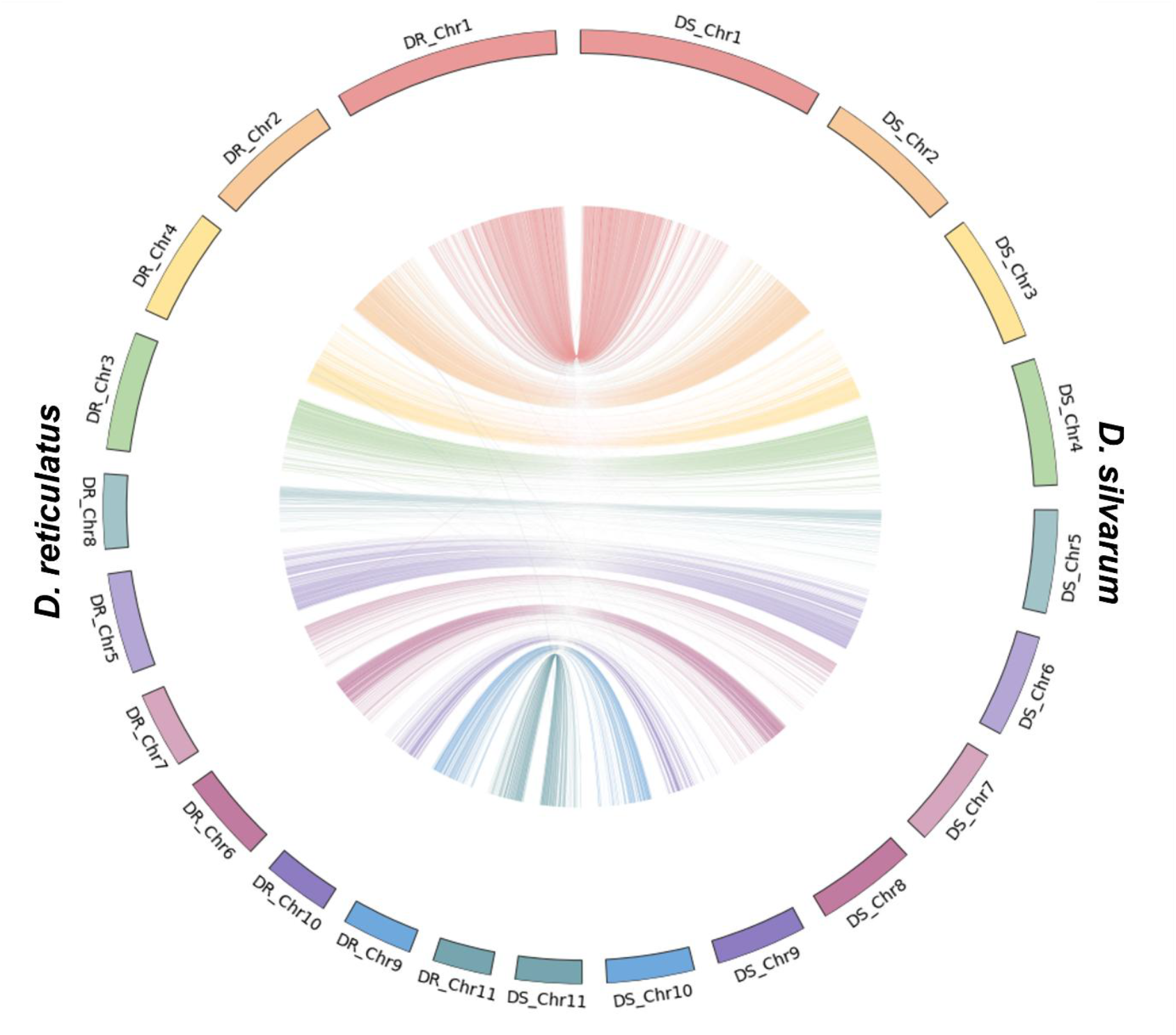
Nucleotide-level Synteny of *Dermacentor reticulatus* and *Dermacentor silvarum* genomes. Nucleotide-level synteny between *Dermacentor reticulatus* (DR) and *Dermacentor silvarum* (DS) is shown for the repeat-masked guided *D. reticulatus* assembly. Scaffolds and links are coloured according to corresponding *D. silvarum* chromosomes.

### Quality of Genome Annotation and Gene Synteny

The unguided and guided *D. reticulatus* genome assemblies were annotated using a combination of homology and *ab-initio* approaches. Prior to annotation, the *D. reticulatus* underwent repeat masking and was found to be repeat-rich, with approximately 61.9% of the genome being comprised of repetitive elements (**Supplementary Figure S1**). Retroelements are the most abundant class, making up 31.85%, and include LINEs (15.63%), LTR elements (15.28%), and SINEs (0.94%). Among LINEs, the R1/LOA/Jockey (5.36%) and RTE/Bov-B (4.67%) subfamilies are particularly prominent. Within LTR elements, Gypsy/DIRS1 elements dominate at 12.12%, indicating a substantial contribution from ancient retroviral-like sequences. DNA transposons account for 2.83%, with Tc1-IS630-Pogo and hobo-Activator being the most common families.Rolling-circle elements contribute a smaller proportion (0.21%). Notably, a large portion of the genome (24.68%) remains unclassified, highlighting potential novel or highly diverged repeat families yet to be characterised. Additional minor components include simple repeats (1.70%), small RNAs (0.47%), satellites (0.09%), and low-complexity regions (0.07%). This composition reflects the dynamic nature of the *D. reticulatus* genome and its evolutionary history, with a predominance of retrotransposon activity and a high degree of uncharacterised repetitive sequence content.

Annotation quality was assessed using BUSCO and compared against other *Dermacentor* assemblies (**Supplementary Table S1**). The *D. reticulatus* assemblies yielded BUSCO completeness scores of 95.0% (unguided) and 95.1% (guided), slightly lower than those of *D. silvarum, D. andersoni*, and *D. albipictus*, all with approximately 98%. Notably, the higher scores in these other species appear to be primarily driven by elevated numbers of complete duplicates (mean = 1160), which are lower in *D. reticulatus* (unguided: 759; guided: 793).

The unguided assembly marginally increased the quality of annotation (+0.1%), suggesting minimal impact of scaffolding on gene model recovery. Furthermore, we also compared gene synteny across the *Dermacentor* assemblies (**Figure 2**). Notably, *D. reticulatus, D. silvarum* and *D. andersoni* all have 11 chromosomes, whilst only 10 are reported for *D. albipictus*. The gene order across the *Dermacentor* assemblies was largely conserved, with minor rearrangements such as on chromosome 1 between *D. reticulatus* and *D. andersoni*. For some chromosomes of *D. andersoni*, including chromosomes 2, 3 and 7, the entire chromosomes appear to be inverted, but the order of genes remains intact (**Figure 2**). This is likely due to the different approaches in the assembly process or interpretation of Hi-C data. *D. reticulatus* had the highest gene synteny with *D. silvarum*, which is expected given their closer phylogenetic relationship and use of the *D. silvarum* assembly to guide scaffolding. The *D. albipictus* genome demonstrates the lowest gene synteny. Chromosome 1 appears to be a combination of inverted (putative) chromosomes 1 and 2 of *D. reticulatus* and *D. silvarum*. Despite these structural differences, gene order along the chromosomes was well-conserved across species, suggesting high fidelity in gene annotation and cross-species genomic alignment.

**Figure 2.**
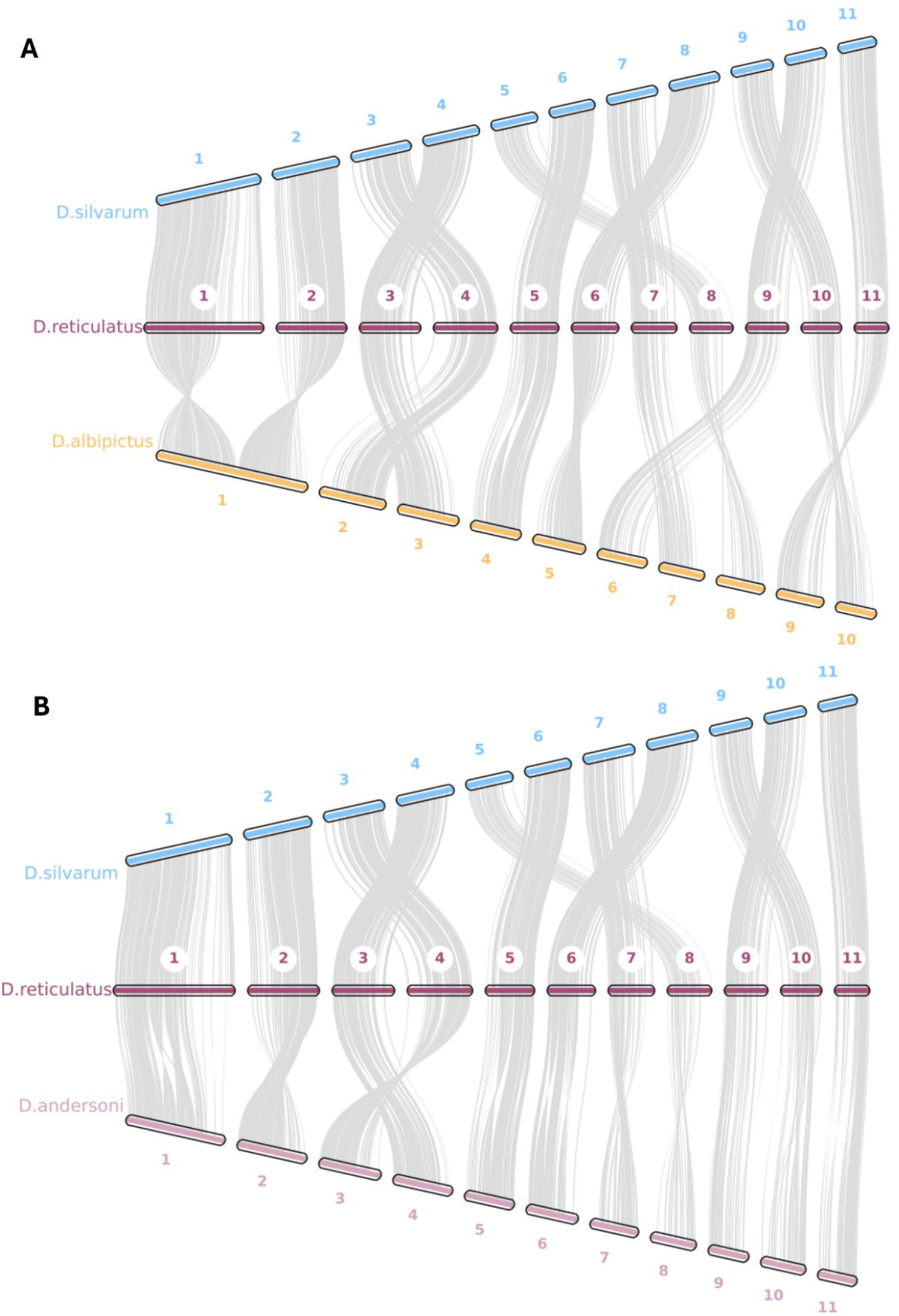
Gene-level synteny between the guided *Dermacentor reticulatus* assembly and members of the *Dermacentor* genus. Gene-level synteny (1:1 orthologous regions) between the guided *D. reticulatus* genome assembly and *D. silvarum* (guiding reference; GCA_013339745.2, [12]). Comparisons are also made to *D. albipictus* (GCA_038994185.2) (A) and *D. andersoni* (GCA_023375885.3) (B). Only synteny between (putative) chromosomes are shown. Blocks are coloured according to each species.

### Mitochondrial Genome Analysis of Tick Species

We also assembled and annotated the mitochondrial genome of *D. reticulatus*, which is 15,103bp in length and contains 38 genes, including 23 tRNAs and 2 rRNAs (**Supplementary Figure S2**). A comparison to 189 publicly available tick mitochondrial genomes, including a previously published *D. reticulatus* mitogenome (NC_068757.1 [51]), confirmed the completeness and accuracy of our assembly, with strong clustering of the two *D. reticulatus* sequences (**Figure 3**).

**Figure 3.**
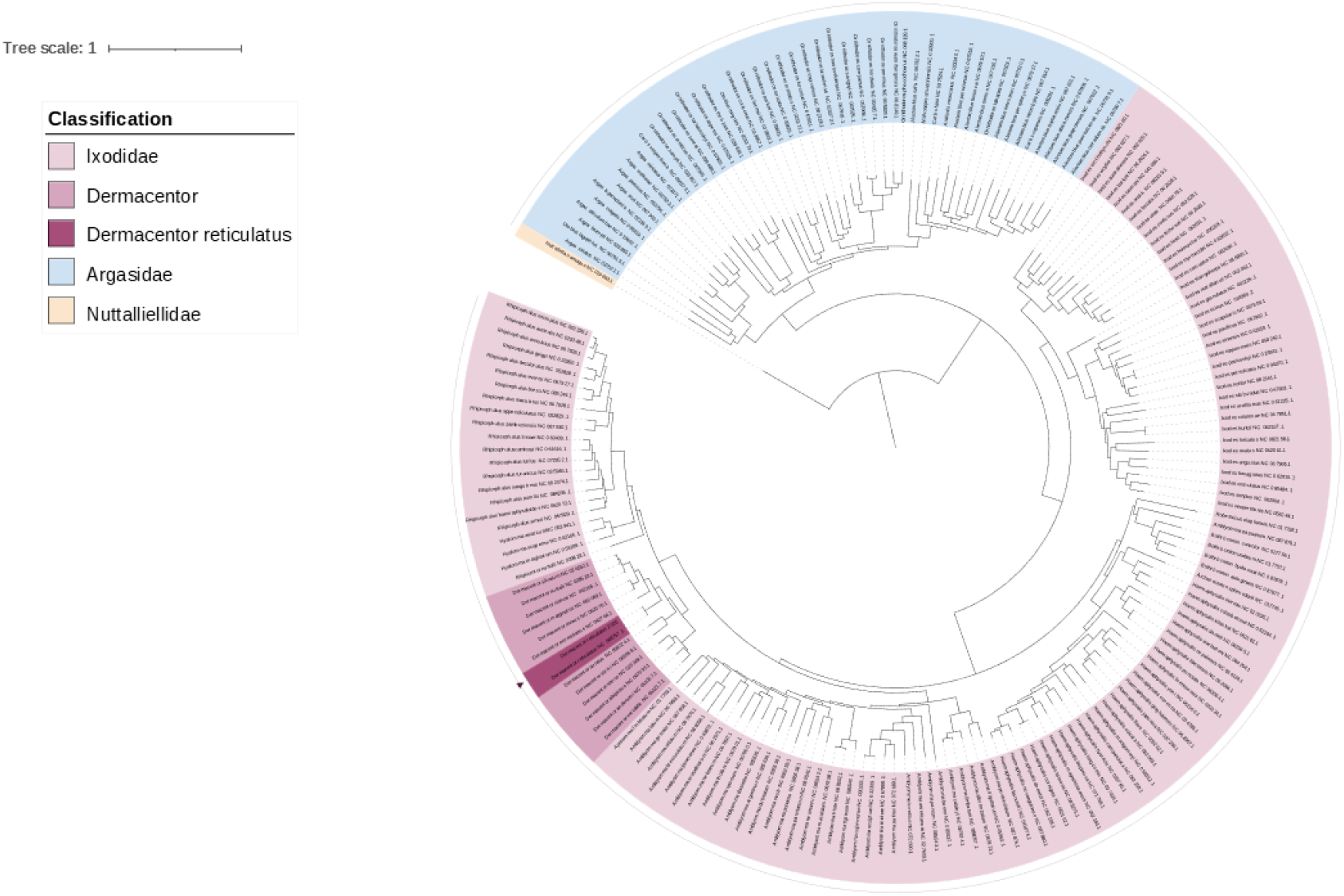
Maximum-likelihood Tree of Tick Mitochondria Sequences. A Maximum-likelihood tree was constructed using an alignment of 189 mitochondria sequences across tick species. The new reference for *D. reticulatus* is highlighted, showing expected clustering within the *Dermacentor* genus. Clades are coloured according to their classification.

Phylogenetic analysis of mitochondrial sequences revealed broader evolutionary relationships across tick families. *Argasidae* (soft ticks) and *Ixodidae* (hard ticks) formed distinct but closely related clades, while *Nuttalliellidae*, represented by one species *Nuttalliella namaqua* from Southern Africa, diverged earlier in the phylogeny. The *Dermacentor* genus is most closely related to the *Rhipicephalus* and *Hyalomma* genera. Within *Dermacentor*, phylogenetic patterns largely reflected geographic distribution. *D. reticulatus* clustered amongst *D. silvarum, D. marginatus, D. nuttalli, D. sinicus, D. niveus*, and *D. everestianus*, suggesting a shared evolutionary history across Europe, Central, and East Asia. *D. auratus* and *D. steini*, found in South and Southeast Asia, also share a close relationship. North American species such as *D. albipictus, D. andersoni*, and *D. variabilis* are less closely related to *D. reticulatus*, consistent with their distinct geographic origins.

### Comparison of Gene Orthologues of Tick Species

We further performed a comparison between annotated genome sequences to gain a comprehensive overview of gene evolution, orthology and duplications (**Figure 4, Supplementary Table S2**). The guided *D. reticulatus* genome assembly contained 52,454 predicted genes, exceeding the gene count observed in other tick species (range: 24,211–46,983). Notably, *D. albipictus* also exhibited a high number of genes (n=46,983), which may explain its high BUSCO score for protein sequences. The elevated number of predicted genes in *D. reticulatus* may in part reflect overprediction by the GALBA annotation pipeline, possibly due to the greater number of scaffolds, which can lead to artificial gene duplications or false positives. Conversely, the lower gene counts observed in other tick assemblies may be attributed to incomplete annotations or underprediction due to differences in assembly quality and annotation methods.

**Figure 4.**
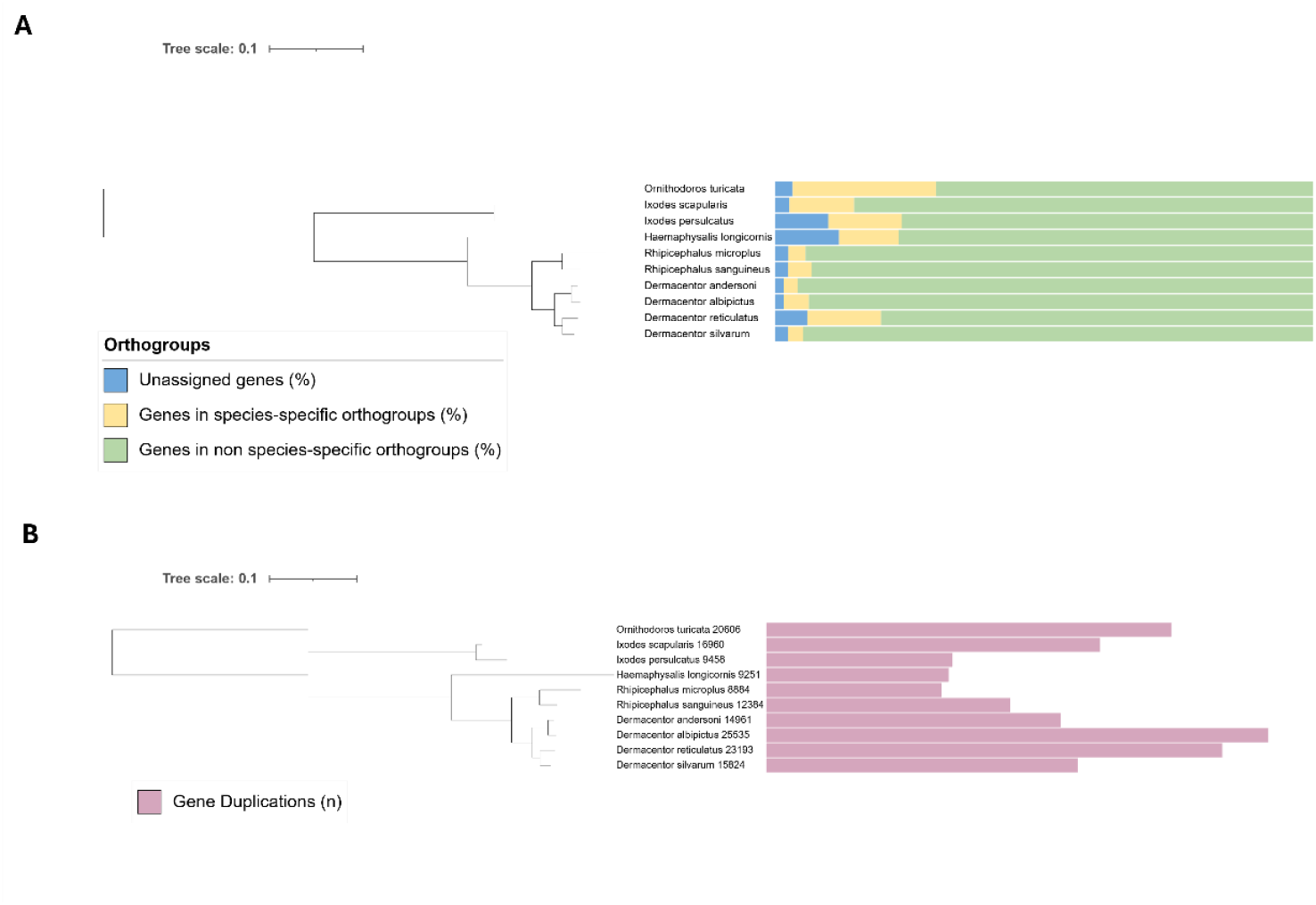
Comparative analysis of Gene Orthologues in Tick species. Maximum-likelihood trees constructed using gene orthologues (A) and duplications (B) across tick species, including *D. albipictus* (GCF_038994185.2), *D. andersoni* (GCA_023375885.3), *D. silvarum* (GCF_013339745.2), *Haemaphysalis longicornis* (GCA_013339765.2), *Ixodes scapularis* (GCF_016920785.2), *Ixodes persulcatus* (GCA_013358835.2), *Ornithodoros turicata* (GCF_037126465.1), *Rhipicephalus microplus* (GCF_013339725.1) and *Rhipicephalus sanguineus* (GCF_013339695.2). Trees are rooted on the *O. turicata* outgroup. Additional comparative genomic statistics are also shown. Figure 3A bars indicate the percentage of genes belonging to non-species-specific orthogroups, species-specific orthogroups and unassigned genes. Figure 3B bars indicate the number of gene duplications and are labelled (left) with the corresponding number of duplications.

Despite this, orthology assignment was successful across species. Approximately 94% of *D. reticulatus* genes were assigned to orthogroups. A maximum-likelihood phylogenetic tree built from single-copy orthologous genes supported expected evolutionary relationships, clustering *Dermacentor* species more closely with other members of the *Ixodidae* family, while *Ornithodoros turicata* (a soft tick from the family *Argasidae*) formed an outgroup (**Figure 4A**).

Different patterns were observed for gene duplication events. A total of 23,193 gene duplication events were observed for *D. reticulatus*, similar to values for *D. albipictus* (n=25,535) (**Figure 4B**). The number of gene duplication events were lower for *D. silvarum* (n=15,824) and for *D. andersoni* (n=14,961). This may also be due to the different completeness and accuracy of annotation approaches used across the *Dermacentor* species.

### Comparative Functional Genomics and GO Term Analysis

Following the analysis of tick evolutionary relationships based on orthologous and mitochondrial sequences, additional insight into key biological adaptations and traits underlying tick evolution was gained through protein family (Pfam) domain comparisons and gene ontology analysis (**Supplementary Tables 3-11**). These approaches have previously been applied in genomic studies of other tick species and provide a broader functional context for interpreting evolutionary changes [12, 53]. In the absence of biological replicates, we compared the fold change of each domain and its associated GO term between members of the *Dermacentor* genus and three other groups: (1) other tick species, (2) blood-feeding insects [BFI: *Aedes aegypti, Anopheles gambiae* and *Aedes albopictus*], and (3) non-blood-feeding arachnids [NBFA: *Centruroides sculpturatus* and *Parasteatoda tepidariorum*] (see **Methods**), similar to Jia *et al*., 2020 [12]. Across all comparisons, most enriched GO terms corresponded to molecular function (Other ticks: 143; BFI:125 and NBFA: 89), followed by biological processes (Other ticks: 107; BFI:88 and NBFA:63) and cellular components (Other ticks: 38; BFI:30 and NBFA:18). These enriched terms covered a wide range of functions, many of which may reflect evolutionary adaptations relevant to hematophagy and vector competence (**Supplementary Tables S3-S11**).

Domains associated with multicellularity were more prevalent in the *Dermacentor* genus compared to other tick species (maximum fold change: 7.25) and BFIs (maximum fold change: 4.83). Among molecular function categories, mannosyl-glycoprotein endo-beta-N-acetylglucosaminidase activity was amongst the highest average fold change across all comparator groups, particularly when compared to other ticks and BFIs (fold change: 13 in both) and NBFAs (fold change 8.67). We also observed a marked enrichment of domains related to metalloendopeptidase activity in *Dermacentor* compared to NBFAs (fold change: 6.37). These enzymes are commonly implicated in extracellular matrix degradation, which could be important in the blood-feeding process. Another key biological process pinpointed when evaluated alongside mosquitoes (BFIs) was cholesterol metabolism (**Supplementary Table S6**). Ticks and mosquitoes cannot synthesise cholesterol *de novo* and must acquire it from external sources. However, mosquitoes can convert plant-derived sterols into cholesterol via modification pathways [54], whereas ticks, being obligate blood feeders, depend entirely on their vertebrate hosts for cholesterol acquisition [55]. The enrichment of cholesterol metabolism-related GO terms in *Dermacentor* may reflect adaptations to this dependency. A minor but notable enrichment in cholesterol-related terms was also detected when comparing *Dermacentor* to other tick species (**Supplementary Table S3)**, suggesting possible lineage-specific metabolic adaptations. Finally, several enriched GO categories across all comparisons were related to nutrient metabolism, including lipid, carbohydrate, glucose, fatty acid, amino acid, mannose, and fructose metabolism, as well as cellular components such as the extracellular matrix (**Supplementary Tables S3–S11**).

Previous studies have also highlighted the importance of metallobiology (heme metabolism), serpins (protease inhibitors involved in immunity and blood feeding), defensins (antimicrobial peptides) and detoxification pathways (cytochrome P450s, glutathione S-transferases and ABC transporters) as key adaptations underpinning tick biology and survival [53]. Therefore, we searched for protein domains associated with these functional categories using InterProScan and identified putative heme biosynthesis proteins in *D. reticulatus* via homology to annotated proteins from the *Dermanyssus gallinae* mite transcriptome [46]. Among the protease inhibitors, we detected 13 proteins with the core cystatin domain (IPR000010) and 43 proteins containing the core serpin domain (IPR023796), indicating active roles for these inhibitors in modulating host immune responses and regulating proteolytic activity during blood feeding (**Supplementary Table S12**). Within the defensin family, only one hit was found for the invertebrate/fungal defensin domain (IPR001542), suggesting a limited but present antimicrobial peptide repertoire (**Supplementary Table S12**). For detoxification-related proteins, we identified 97 cytochrome P450 domain-containing proteins (IPR001128), 77 carboxylesterases (IPR002018), and multiple glutathione S-transferases (22 hits for the N-terminal domain [IPR004045] and 27 for the C-terminal domain [IPR004046]). Additionally, 134 proteins contained ABC transporter-like domains (IPR003439) and 33 had transmembrane domains (IPR011527), suggesting the expansion of ABC transporter-mediated efflux mechanisms (**Supplementary Table S12**). These domain profiles support the presence of complex and specialised molecular systems in *D. reticulatus* for managing oxidative stress, host interactions and xenobiotic detoxification [20, 56-59].

To investigate the heme biosynthesis pathway in *D. reticulatus*, we searched for homologs of key enzymes using *Dermanyssus gallinae* proteins as references. Homologs for the late pathway enzymes, including protoporphyrinogen oxidase (PPOX) and ferrochelatase (FECH), were identified (1.31E-133 and ≤8.09E-136, respectively), indicating conservation of the terminal steps of heme biosynthesis (**Supplementary Table S13)**. However, no matches were detected for several early enzymes in the pathway, such as 5-aminolevulinate synthase (ALAS1/ALAS2), ALA dehydratase (ALAD), porphobilinogen deaminase (HMBS), uroporphyrinogen III synthase (UROS), and uroporphyrinogen decarboxylase (UROD), suggesting these genes may be absent. This finding aligns with previous reports indicating that Metastriata ticks have lost enzymes involved in the early stages of the pathway [53].

## Discussion

*Dermacentor reticulatus* is a carrier of several tick-borne pathogens, including *Babesia, Borrelia, Anaplasma, Rickettsia, Bartonella, Francisella, Coxiella* and tick-borne encephalitis virus [6]. Due to anthropogenic and climate factors, *D. reticulatus* has expanded its range to regions previously uninhabited by this tick species [3]. Consequently, effective surveillance of *D. reticulatus* is crucial for managing the risk of transmitting tick-borne diseases that pose substantial medical and veterinary threats. Such surveillance schemes include the UKHSA Tick Surveillance Scheme (TSS) that has been implemented in the United Kingdom [4, 60]. Since 2024, *D. reticulatus* has been reported using TSS in known endemic locations (e.g. Wales, Essex and South West England) [5]. While some of the ecology and behaviour of *D. reticulatus* is better understood, its genome remains largely unexplored and no chromosome-scale genome assembly is currently available. Genomes of other tick species, such as *Ixodes scapularis* (the black-legged tick), have been sequenced and annotated in detail [12, 38]. There remains a limited understanding of the unique molecular and biological adaptations in *D. reticulatus*, which might differ significantly from other ticks. Therefore, a high-quality reference genome for *D. reticulatus* will provide a foundation for downstream comparative genomic analysis, enabling more detailed comparisons with other tick species and potentially revealing novel genes or pathways that could be targeted for tick surveillance or disease prevention.

We have shown the genome assemblies of *D. reticulatus* are of comparable quality to existing reference genomes within the *Dermacentor* genus. We included both unguided and guided assemblies, along with their annotations, to capture the structural accuracy and improved contiguity provided by the guided assembly, while also acknowledging that reference-based guidance may introduce bias. [61]. Our analysis reveals that, despite differences in chromosome orientation and potentially chromosome number across the *Dermacentor* genus, there is conservation of nucleotide sequences and gene synteny among these species. Subtle differences may be observed due to differences in the assembly approach used. For example, unlike previous studies, we utilised long-read ONT data to generate the draft assembly. Incorporating Hi-C and Illumina data could have further improved the assembly by enhancing chromosomal-level scaffolding and correcting potential structural variations [20]. However, we demonstrate the utility of ONT data for rapid, cost-effective genome assembly of a non-model organism using high molecular weight DNA, especially for complex organisms, including ticks. The use of ONT data offers significant advantages for assembling large, repetitive genomes like that of *D. reticulatus*, which we estimate to be approximately 60% repetitive [62]. As a result, we have generated both unguided and guided assemblies of *D. reticulatus*, each approximately 2.3 Gb in size, achieving high BUSCO scores (>96%), indicating strong completeness and accuracy in the genome reconstruction.

Additionally, we annotated our genome assemblies using both homology-based and *ab initio* approaches. Our predictions indicate that *D. reticulatus* contains 58,384 genes, but notably exhibits fewer complete duplicated genes compared to other species within the *Dermacentor* genus. This observation could potentially be refined with the incorporation of RNA-seq data and improved contiguity, which would enhance gene model accuracy and provide a clearer understanding of gene annotation. Furthermore, we performed an orthologous gene analysis to explore the evolutionary relationships and functional similarities of genes in *D. reticulatus* compared to other tick species, including *D. albipictus, D. andersoni, D. silvarum, H. longicornis, I. scapularis, I. persulcatus, O. turicata, R. microplus*, and *R. sanguineus*. Our analysis involved identifying orthologous genes across these species, providing insights into their shared ancestry and functional conservation. This work identified a significant number of orthologous gene clusters across all species, suggesting that while ticks diverged along different evolutionary paths, many genes have been conserved throughout the Ixodida order. These conserved genes are likely to play fundamental roles in basic cellular processes, such as metabolism, DNA repair and immune response, which are critical for tick survival and adaptation to various environmental niches and hosts [20, 53]. We also observed differences in the number of duplicated genes between *D. reticulatus* and other members of the *Dermacentor* genus, which may have contributed to species diversification. However, further validation of these gene predictions using RNA-seq data is needed to better characterise gene orthology, duplication events, and their potential functional significance.

Our functional domain and GO enrichment analyses shed light on several molecular adaptations in *D. reticulatus* that support its hematophagous lifestyle [20]. The identification of detoxification-related domains, including cytochrome P450s, carboxylesterases, glutathione S-transferases, and ABC transporters, suggests a capacity to process host-derived and environmental xenobiotics, as well as to mitigate oxidative stress associated with blood feeding [56, 57]. Similarly, the presence of abundant serpin and cystatin domains highlights tight protease regulation, likely crucial for controlling host responses and maintaining gut homeostasis during prolonged feeding [63]. Although defensin-related domains were limited, this does not preclude the existence of alternative antimicrobial strategies [58, 59].

Importantly, we observed strong enrichment of metalloendopeptidase-related GO terms in *Dermacentor* compared to non-blood-feeding arachnids. These enzymes, often involved in extracellular matrix degradation, may facilitate host tissue penetration or remodelling during tick attachment and feeding. For example, thiol-activated metalloendopeptidase with kininase activity, a protease enzyme that breaks down peptides, has been observed in *Boophilus microplus* saliva (cattle tick) [64]. The enrichment of cholesterol metabolism-related GO terms in *Dermacentor* was also of interest. As obligate blood feeders, ticks cannot synthesise cholesterol and must acquire it entirely from their vertebrate hosts. In contrast, mosquitoes retain the ability to modify plant sterols, reducing their dependence on host cholesterol [54]. The overrepresentation of cholesterol-associated pathways in *D. reticulatus* may therefore represent enhanced systems for uptake, trafficking or utilisation of host-derived lipids. Further, while late-stage heme biosynthetic enzymes (PPOX and FECH) were present in *D. reticulatus*, earlier pathway components were not detected, consistent with the hypothesis that Metastriata ticks have lost the ability to synthesise heme. This suggests a reliance on host-derived heme, as previously observed for *D. silvarum* and other tick genera [12, 53].

Finally, we present the mitochondrial genome of *D. reticulatus*, along with a phylogenetic tree constructed from mitochondrial data available across all tick species. The mitochondrial genome has a rapid evolutionary rate, is maternally inherited, and has a relatively conserved structure across species. As such, mitochondrial markers are important for inferring evolutionary relationships, especially among closely related species within a genus or across genera [65]. The mitochondrial maximum-likelihood tree we constructed clarifies the evolutionary position of *D. reticulatus* within the *Dermacentor* genus and reveals notable geographic patterns, suggesting the influence of geographic barriers or migration routes on its distribution. Given the high conservation of the mitochondrial genome in *D. reticulatus*, additional nuclear genomic markers will be essential for identifying signatures of geographical adaptation within this species.

## Conclusion

In summary, we have generated the first draft reference genome for *D. reticulatus* using ONT long read data and provided a mitochondrial sequence alongside a phylogenetic tree, which reveals important geographic patterns within the genus. The annotated genome offers insights into key genes and biological processes, establishing a valuable resource for future population genetic studies and for advancing our understanding of tick evolution, one-health and infection control.

## Abbreviations

(ONT): Oxford Nanopore Technologies
(DR): *Dermacentor reticulatus*
(DS): *Dermacentor silvarum*
(BUSCO): Benchmarking Universal Single-Copy Orthologs
(BFI): blood-feeding insects
(NBFA): non-blood-feeding arachnids

## Declarations

NA

## Ethics approval and consent to participate

NA

## Consent for publication

NA

## Availability of data and materials

All raw sequencing data have been submitted and genome assemblies and annotations have been submitted to NCBI (Accessions TBC) and LSHTM genomics data repository: https://genomics.lshtm.ac.uk/data/.

## Authors’ contributions

SMB and FVB coordinated and delivered fieldwork, including tick collection and morphological species identification. EM, MSJ, SC, and MK carried out laboratory work, including sample processing, high molecular weight DNA extraction and Oxford Nanopore Technologies (ONT) sequencing. NB performed the bioinformatic analyses and result interpretation with assistance from JT and under the supervision of TGC and SC. NB drafted the manuscript, with input and critical revision from SC, TGC and other co-authors. All authors contributed to discussions throughout the study, reviewed and edited manuscript drafts, and approved the final version for publication.

