## Supplementary Figure for "The First Chromosome-Scale Genome Assembly for *Dermacentor reticulatus*: A Key Vector of Tick-Borne Pathogens of Public and Veterinary Health Importance in Europe"

**Supplementary Figures**

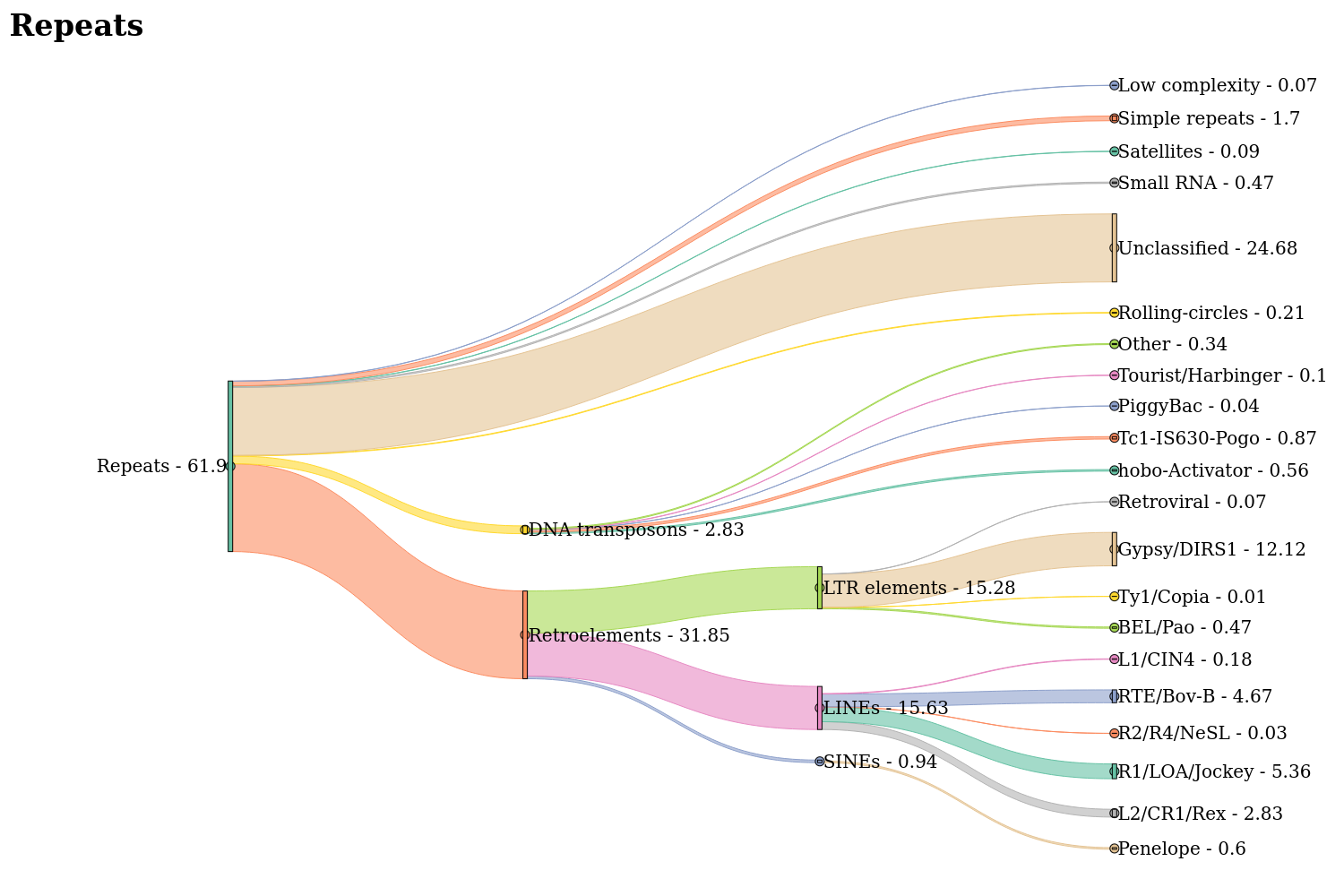

**Supplementary Figure S1. *Repeat content of the* Dermacentor reticulatus *genome visualised as a Sankey diagram.***
The figure illustrates the hierarchical classification and relative abundance of repetitive elements identified in the *D. reticulatus* genome. Major repeat classes are shown as primary branches, including retroelements (31.85%), DNA transposons (2.83%), rolling-circle elements (0.21%), unclassified repeats (24.68%), and other categories such as small RNAs, satellites, simple repeats, and low-complexity regions. Retroelements are further subdivided into SINEs (0.94%), LINEs (15.63%), and LTR elements (15.28%), with their respective subfamilies (e.g., R1/LOA/Jockey, Gypsy/DIRS1, BEL/Pao) shown according to their genomic proportions. Similarly, DNA transposons are broken down into subtypes such as hobo-Activator, Tc1-IS630-Pogo, and others. The width of each flow is proportional to the genomic percentage of the corresponding repeat category. This visualization highlights the dominance of retroelements, particularly LINEs and LTR elements, as well as the substantial proportion of unclassified repeats in the *D. reticulatus* genome, reflecting its repeat-rich architecture.

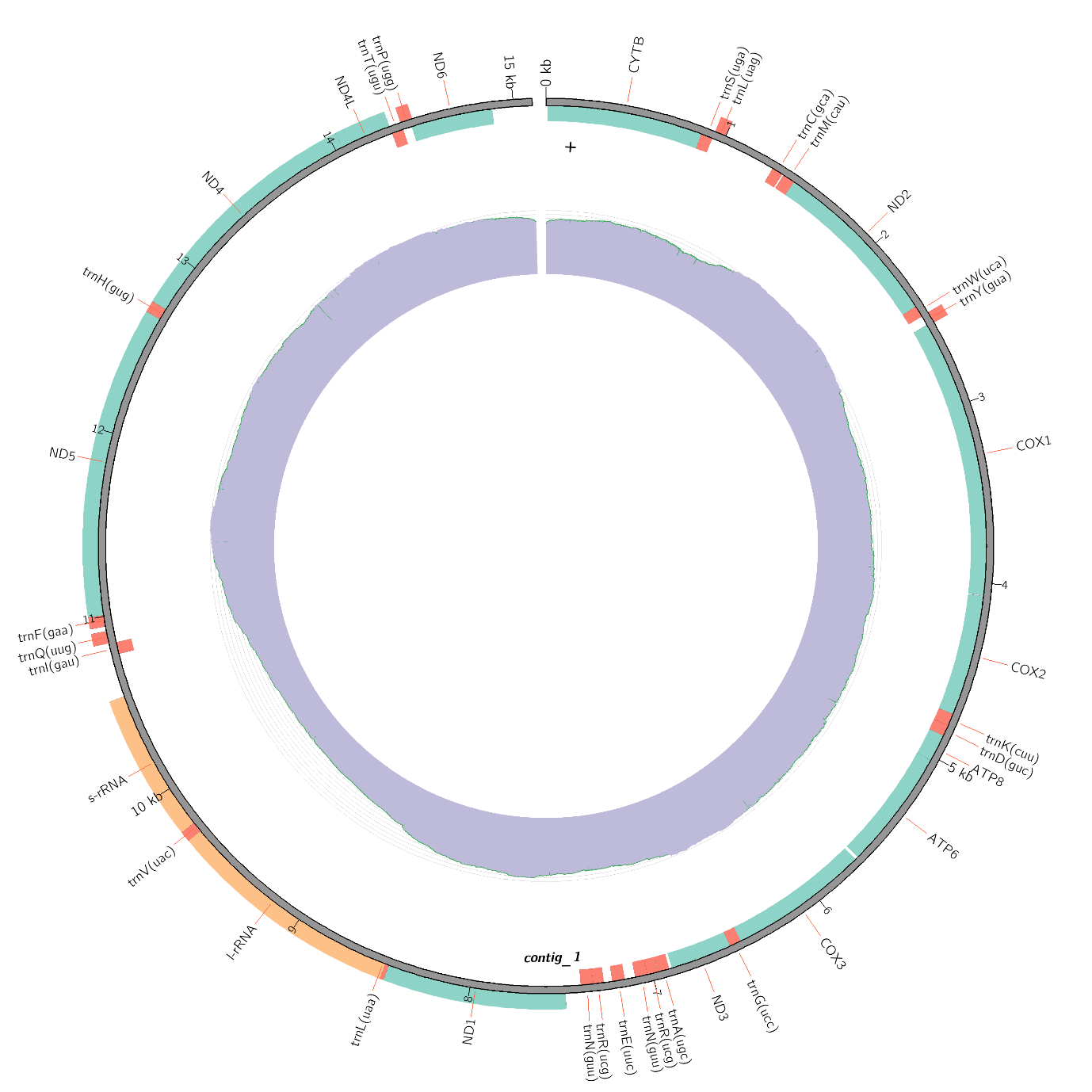

**Supplementary Figure S2. Mitochondria Sequence and Annotation for *D. reticulatus.*** The circular mitochondria sequence for *D. reticulatus* is shown alongside its annotation and genome coverage.
